# CryoSift – An accessible and automated CNN-driven tool for cryo-EM 2D class selection

**DOI:** 10.1101/2025.07.28.667259

**Authors:** Jan-Hannes Schäfer, Austin Calza, Keenan Hom, Puneeth Damodar, Ruizhi Peng, Nebojša Bogdanović, Gabriel C. Lander, Scott M. Stagg, Michael A. Cianfrocco

## Abstract

Single-particle cryo-electron microscopy (cryo-EM) has become an essential tool in structural biology. However, automating repetitive tasks remains an ongoing challenge in cryo-EM dataset processing. Here, we present a platform-independent convolutional neural network (CNN) tool for assessing the quality of 2D averages to enable automatic selection of suitable particles for high-resolution reconstructions, termed CryoSift. We integrate CryoSift into a fully automated processing pipeline using the existing cryosparc-tools library. Our integrated and customizable 2D assessment workflow enables high-throughput processing that accommodates experienced to novice cryo-EM users.

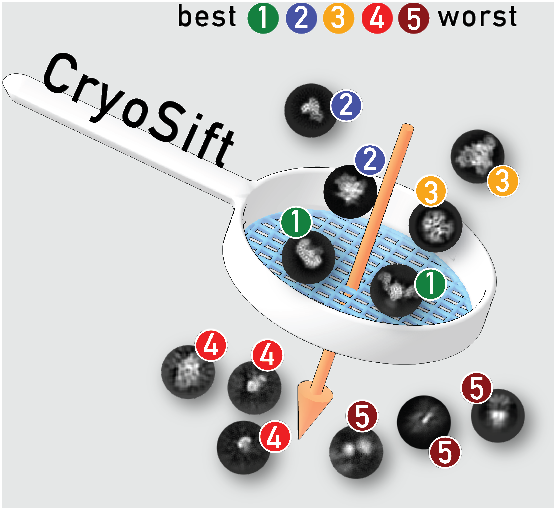

## Introduction

Single-particle cryogenic electron microscopy (cryo-EM) has rapidly grown from a niche method into a fundamental tool in structural biology ^1^. While this evolution has made processing of challenging macromolecules more accessible, the cryo-EM community would benefit from reliable automated processing workflows. Current single-particle analysis (SPA) applications in cryo-EM are required to be reliable, fast, and user-friendly. The widespread adoption of cryo-EM has been facilitated by graphical user interfaces. Thus, new automated pipelines must not only be accessible via the command line but also incorporated into user-friendly interfaces.

Many steps in preprocessing raw cryo-EM data to particle stacks and 2D class averages are nearly automated. The stack of sequential frames that comprise a single cryo-EM micrograph must first be aligned to account for electron beam-induced motions during acquisition, then aligned frames subsequently averaged so that the aberrations associated with the contrast transfer function (CTF) can be estimated. These processing steps are performed on each of the thousands of images that are associated with a given single particle cryo-EM dataset. Live and streamlined preprocessing tools have been implemented in popular processing environments such as cryoSPARC ^2^ and RELION ^3^, allowing users to adjust data collection according to their specifications. However, the subsequent detection, extraction, and classification of particles remains an iterative process that typically requires user intervention. Since cryo-EM produces high-resolution reconstructions from noisy, heterogeneous particle collections, classifying individual 2D projections of particles into homogeneous groups is often non-trivial.

Automated workflows help address two different bottlenecks. First, there are new users who need guidance at the decision points, such as deciding what are “good” and “bad” images or class averages. Tools that provide assessment metrics are useful for guiding such users to make good decisions for their structure determination, overcoming a “user-input bias.” Second, for well-established samples where investigators are looking at variations, cofactors, drugs bound, and more, it is useful to have tools that take the manual decisionmaking steps out of the workflow so that multiple samples can be processed systematically, as shown for implementing routine processing of G protein-coupled receptors ^4^.

Substantial progress has been made toward a more user-friendly and unsupervised selection of suitable 2D projections. RELION’s Class Ranker automates 2D class selection using a trained convolutional neural net-work (CNN) that labels each class and applies a cut-off, effectively ranking and selecting suitable classes for downstream analysis ^3^. Similarly, SPHIRE ^5^ has implemented a CNN-based labeling approach termed Cinderella (https://github.com/MPI-Dortmund/sphire_classes_autoselect), which separates 2D classes into good and bad class averages. Cinderella offers both a pre-trained CNN model and training options for user datasets. While these methods utilize CNNs, cryoSPARC offers its own approach for automating 2D class selection through similarity comparisons to projections of a user-provided 3D volume. This approach, currently under active development, is expected to be most valuable for processing pipelines of known targets, using shape-similarity and pixel-by-pixel similarity to calculate projections of the provided 3D volume. Our own previous work on removing subjective decision-making for 2D class selection as part of a progressively user-free processing pipeline ^6^ also relies on a CNN-based classifier that automatically selects suitable 2D class averages. This classifier (2DAssess) was trained on a labeled dataset of 2D class averages, preprocessed and categorized into four classes: good, clip, edge, and noise. While 2DAssess showed promise, it was command line only and not incorporated into any user-friendly workflows, limiting adoption.

Current approaches for automated 2D class selection are either platform-dependent or stand-alone command-line tools that do not readily support cross-platform integration. In this study, we addressed this limitation by developing CryoSift, a 2D class average quality assessor that is platform agnostic (i.e., able to assess data from RELION or cryoSPARC) to enable unsupervised dataset processing. We show the utility of this tool by incorporating it into cryoSPARC using the cryosparc-tools API, providing customizability for advanced users while maintaining ease of use for cryo-EM beginners utilizing the default cryoSPARC implementation.

## Material & Methods

### Training of the 2D assessor CryoSift

#### Generating 2D averages for training

EMPIAR datasets from **Table 1** were imported into cryoSPARC (v.4), preprocessed with CTFFIND4 and patch-based motion correction for imported movies. Templates for template-picking were created from associated EMDB maps **Table 1** using cryoSPARC’s ‘Create Templates’ job. Particle dimensions from measurements in ChimeraX ^7^ were used in template picking. Picked particles were extracted using box sizes about 1.5-fold the particle diameter and binned by two. The extracted particles were subjected to two or three rounds of 2D classification. To cover a broad range of 2D class qualities, the projections from all iterations were used for labelling. Additionally, pre-labeled 2D averages from RELION’s 2D Class Ranker (EMPIAR-10812) were utilized. The class averages and selected metadata including the pixel size, resolution estimates from Fourier Ring Correlation (FRC) and relative class distribution were extracted from the processing runs and were presented to the CNN for training.

**Table 1.** Publicly available datasets used to train CryoSift. Datasets with ^*^ were also used for the mass estimator.

| Target | EMPIAR | EMDB | Target | EMPIAR | EMDB | Target | EMPIAR | EMDB |
| --- | --- | --- | --- | --- | --- | --- | --- | --- |
| E3 Ub ligase | 11501 | 16087 | Pol-SN | 11522 | 28786 | Urease | 10389 | 10835* |
| Nucleosome | 11840 | 16859* | Primase | 11131 | 26346 | bGal | 10061 | 2984* |
| E3 Ub ligase | 11734 | 41996 | PreP | 10937 | 22278 | RNApol | 10709 | 12885* |
| Antibody | 11341 | 28537* | ALK | 10930 | 24095 | PA3488 | 11204 | 32438* |
| AP2 Tgn38 | 11605 | 24712* | CST | 10718 | 21567 | GS-GN | 11139 | 14587* |
| AP2 Heparin | 11604 | 24711 | RNAPol-III | 10168 | 4180 | FAsyn | 10470 | 4578* |
| Pks13 | 11608 | 26574 | RAG1-2 | 10049 | 6487 | Cas7-11 | 11268 | 14848* |
| MSG | 11167 | 34029 | N2ase | 11795 | 26764* | T20S | 10025 | 6287* |
| Aldolase | 10379 | 21492* | PRex-1G | 10285 | 20308* | CDK | 10561 | 12042 |
| MAVS | 10031 | 6428 | HA-trimer | 10097 | 8731 | 7TM-1G | 10288 | 0339 |

**Table 2.** Laboratory datasets used for training CryoSift.

| Target | Comment | Target | Comment | Target | Comment |
| --- | --- | --- | --- | --- | --- |
| Tubulin | heterodimer | RNA | 80 kDa | Kinesin binder | complex |
| Protein | 70 kDa | RNase P | RNA-free | HIV-1 | hexamer |
| Spike | SARS-CoV2 | GPCR | complex | 40S | Ribosome |
| Aldolase | EMD-43528 | PolQ | Polymerase | PolQHel | Polymerase |
| PolQHelNt | Polymerase | LONP1 | Protease | NPM1 | pentamer |
| CRBN-DDB1 | Ligase | TTR | tetramer |  |  |

**Table 3.** Test dataset parameters for 2D class average assessment in *CryoSift*.

| Target | EMPIAR | PDB | Images | keV | Pixel size (Å/px) | Particle diameter (Å) |
| --- | --- | --- | --- | --- | --- | --- |
| Aldolase | 10379 | 6ALD | 1,118 | 200 | 0.91 | 100 |
| DPS | 11792 | 6GCM | 194 | 300 | 0.834 | 100 |
| Streptavidin | 10641 | 7DY0 | 2,273 | 300 | 0.40 | 60 |
| 70S Ribosome | 11524 | 8BUU | 6,384 | 300 | 0.82 | 270 |
| P-Rex1G | 10285 | 6PCV | 1,734 | 300 | 1.0 | 70 |
| ApoF | 12798 | 7VD8 | 277 | 200 | 0.74 | 130 |
| RNAPol | 11522 | 8F1K | 6,088 | 300 | 1.083 | 190 |
| Complex-I | 11656 | 7B93 | 571 | 300 | 1.055 | 290 |
| MSG | 11167 | 7YQM | 700 | 300 | 0.822 | 90 |
| Pks13 | 11608 | 8CUZ | 100 | 300 | 0.835 | 170 |

#### Generating mass metadata for CNN training and assessment

Masses for each class average were estimated based on the pixel intensities and pixel sizes. The sum of the pixel intensity of the particle signal in class averages was calculated as follows: 1) Define all pixels 3 standard deviations above the mean as the particle signal and the remainer as background. 2) Subtract the mean of the background from the mean of the particle. 3) Sum the intensity values for the particle pixels and multiply by the pixel size squared. A calibration curve for converting the summed intensities to masses was determined by taking EMPIAR data of homogeneous particles (**Table 1, Fig. S1**) with known masses and plotting the pixel intensities vs their known masses. This resulted in a linear plot, and the slope was used to determine a calibration factor that was multiplied by the summed intensity values for each class average to convert them into kDa masses. A plot of the particles with known masses over the estimated masses for their class averages is shown in **Fig. S1**. The mass information was incorporated into the CNN by including the deviations from the mean, median, and mode mass as part of the metadata used during training and evaluation.

#### Expert labeling of 2D averages for the CNN training

Manual labelling of 2D class averages for training was implemented using the python-based image categorization tool *tkteach* (https://github.com/rmones/tkteach). 2D class averages from cryoSPARC were extracted, converted to JPGs and used as input for an adopted version of *tkteach*. The tkteach GUI allows the easy labeling of the classes (**Fig. 1A**) and storage of the associated labels for training of the CNN. Researchers from the Stagg, Cianfrocco, and Lander labs were involved in labelling the 2D class images. Assessors were provided a rubric for manually grading class averages as follows: A – Best, secondary structure, very sharp looking, B – Decent, secondary structure, some fuzzy regions, C – Acceptable, overall shape of a particle, some domain details, little to no secondary structure, D – Poor, particle-like shape but fuzzy with artifacts, F – Unusable, nothing resembling a particle, artifacts in the background. Assessors were provided with example labelled averages to aid in making consistent evaluations (**Fig. S2**). The pooled averages were used for training the CNN.

**Figure 1.**
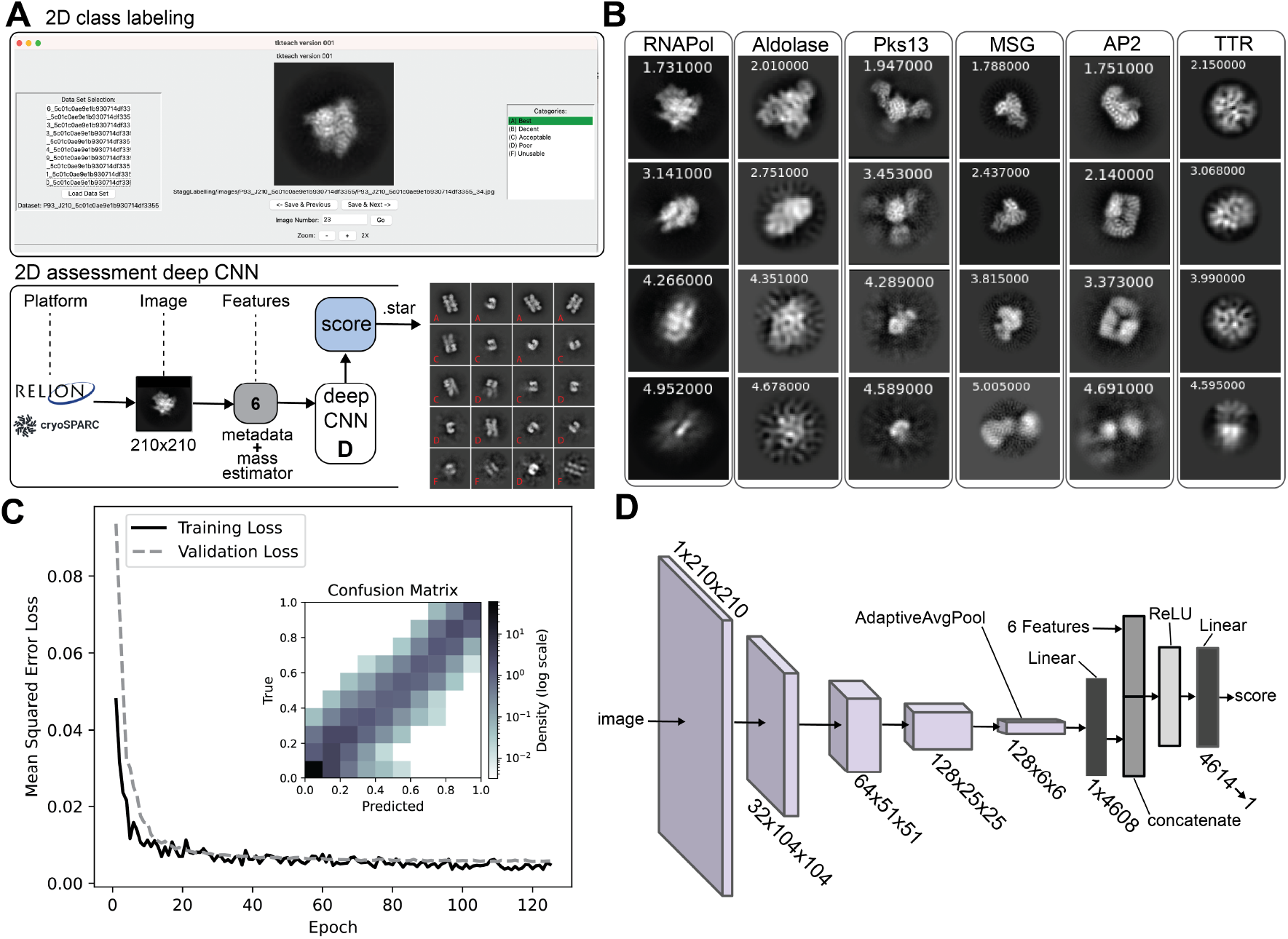
Architecture, training, and benchmarking of CryoSift neural network. (**A**) GUI of *tkteach* for 2D class labeling and basic architecture of the deep convolutional neural network of CryoSift. RELION or cryoSPARC 2D projections of 31×31 to 210×210 px input. Three data features from the mass estimator and 3 metadata features (FRC resolution, class distribution and pixel size) are also fed into the model, resulting in a predicted quality score. Example output of AP2 averages with grade-based labels (red). (**B**) 2D averages with predicted quality scores are grouped by protein, sampled across the full range of class quality scores. (**C**) Mean-square error loss over Epochs of training and validation with features. Insert shows the prediction error between true and predicted score as confusion matrix (density in log scale). (**D**) Overview of the CNN layers (details in **Fig. S3**).

### Designing and training the CNN

Our 2D class average assessor CryoSift was built using residual connections as described in the original ResNet paper ^8^, which allows outputs from earlier layers to bypass one or more subsequent layers, helping to preserve gradient flow during backpropagation and mitigating the vanishing gradient problem caused by repeated application of the chain rule. CryoSift applies two consecutive convolutional filters, each with a 2×2 filter and a stride of 2 instead of pooling. This approach achieves downsampling while also enabling the network to learn weighted combinations of features at each stage, effectively functioning as a form of weighted average pooling. The CNN also employs batch normalization, which takes an input tensor and tries to normalize it to a mean of 0 and standard deviation of 1, averaged across all observed images in the training set. Finally, the CNN employs adaptive average pooling to allows for input images of any size greater than 31×31. This was enabled through the use of PyTorch’s AdaptiveAvgPool2D, which is a pooling layer that adaptively chooses pooling size and stride according to the image’s size, such that the output size of the pooling layer is always the same, in this case 6×6. CryoSift was trained using Python 3 and PyTorch. The Adam optimizer ^9^ was used for model generalization and convergence. The learning rate was 0.0001 with weight decay of 0.0001. The batch size was 32, and the model was trained for 200 epochs with little change to validation loss after approximately 60 epochs.

### Testing and validation of CryoSift

For testing the Cryosuft, 10 single-particle cryo-EM datasets were downloaded from the EMPIAR repository, covering a wide range of molecular weights. Those include Aldolase, DNA protection during starvation protein (DPS), Streptavidin, 70S ribosome in complex with VmlR2, P-Rex1–G-beta-gamma signaling scaffold (P-Rex1G), RNA polymerase Sigma N (RNAPol), Malat synthase G (MSG), mycobacterial polyketide synthase 13 (Pks13), Mitochondrial Respiratory Complex I (Complex-I) and an in-house dataset of mouse apoferritin. The associated identifiers and data collection parameters are listed in **Table 2**.

### Sample preparation and cryo-EM of mouse apoferritin

Mouse apoferritin (heavy chain) was prepared, following our published protocol ^10^, yielding a concentrated sample of 15 mg/ml in 30 mM HEPES pH 7.5, 150 mM NaCl, 1 mM DTT with 5% (v/v). Graphene grids were prepared, following our established protocol ^11^. In brief, methyl-methacrylate-supported graphene was transferred on Quantifoil UltrAuFoil 0.6/1.0 400 mesh grids and treated with ozone to render the graphene hydrophilic. Apoferritin was diluted to 1.5 mg/ml in glycerol-free buffer and applied to the grids and plunge frozen manually at 4 °C at 90% humidity for 3 seconds. 900 movies were collected on a 200 keV Arctica (Thermo Fisher) equipped with a Falcon 4 (Thermo Fisher) at a total electron exposure of 40 e^−^/Å ^2^ and a nominal magnification of 190,000 and an uncalibrated pixel size of 0.74 Å /pixel. Movies were collected automatically using EPU (v.3.9).

### Standardized processing of single particle datasets in cryoSPARC

All datasets from **Table 1** were processed in cryoSPARC (v.4.6), following a standardized processing scheme: If required, beam-induced motion in the imported EMPIAR datasets was corrected using Patch-Motion correction, followed by CTF estimation with Patch-CTF. For the in-house ApoF dataset, micrographs were curated, using a CTF cutoff at 4 Å and astigmatism of lower than 600 Å , yielding a stack of 277 micrographs. Picking-templates were generated from a small stack of 50 micrographs using a blob-picker with the diameters from **Table 1** . The subsequently template-picked particles were extracted in a box matching twice the particle diameter, Fourier-cropped to 100 pixels (px). The resulting 2D classification was used as input for the automated processing pipeline using our cryoSPARC tools implementation with mask sizes 20% larger than the particle diameter from **Table 1**. Non-default parameters were automatically selected from the user-provided extraction box. Box sizes above 300 px are considered large, while boxes below 200 px are considered small. All in-betweens are prompting the default settings. All particles were classified using a 100 px Fourier-cropped box, allowing adaptive binning for larger boxes. Small inputs are processed with the new 2D Classification (Small Particle) job type, using 3 Å maximum resolution and initial class uncertainty factor 3, turned Force max over poses/shifts off, over 40 online-EM iterations with a Batchsize of 400. Ab initio reconstructions were generated using a maximum resolution set to 3 Å , initial minibatch size 300 and final minibatch size 1000. Re-processing of the Pks13, DPS and Complex-I datasets in RELION5 was performed using pre-processed particle stacks from cryoSPARC. For file-conversion and transfer to RELION5, *pyem* (https://github.com/asarnow/pyem/) was utilized. 2D classification in RELION5 was performed with VDAM-enabled, T=2 and 20 classes.

## Results

### Design of the CNN-based 2D assessor CryoSift

Building on RELION’s *Class Ranker* ^3^ and our previous *2DAssess* tool ^6^, we developed a 2D assessment tool that employs a deep CNN to evaluate 2D class averages and their platform-independent metadata, termed *CryoSift*. The model incorporates pixel size, resolution estimates from the FRC, relative class distribution, deviation from mean mass, deviation from median mass and deviation from mode mass as input features **Fig. 1A** . Images with higher class distribution, better resolution, and smaller pixel size typically produce sharper, higher-quality projections, enabling the CNN to leverage these parameters for improved classification. Based on the provided images and associated metadata, the trained CNN assigns continuous quality scores ranging from 1.0 (best) to 5.0 (worst). The model was trained using a mean-square-error (MSE) loss function, not a Softmax-based function (which would limit the score to a certain range). This means the model is technically able to predict real numbers as the score, relative to its training scores range. This results in open score boundaries, and for cases of classes worse than the worst training class, scores higher than 5 are given. For classes better than the best training class, scores smaller than 1 are assigned. The 2D class evaluator accepts images from both RELION and cryoSPARC platforms, enabling platform-independent data processing with cross-platform compatibility.

Our algorithm automatically Fourier-scaled all input images to 210×210 pixels to allow adaptive feature extraction. Images smaller than this dimension are zero-padded to 210 pixels, while larger images are down-sampled. This approach preserves native image features to enhance prediction accuracy, with a minimum input size requirement of 31×31 pixels for zero-padding operations. Padding each 2D class image to a fixed size instead of allowing variable inputs during training results in two main advantages: 1) Allows us to train in mini-batches; in the sense that the model updates its weights after seeing a batch of 32 images, instead of after every image. This allows a much smoother weight convergence, plus leverages GPU parallel hardware to speed up training. 2) Allows us to use batch normalization, a common neural network method that greatly speeds up training convergence. Batch normalization requires the model to train on mini-batches of data.

The architecture of our deep CNN is detailed in **Fig. 1D** (layer details in **Fig. S3**). The CNN was trained on 32,204 total 2D averages: 26,389 pre-labeled images from EMPIAR-10812 (used in RELION’s *Class Ranker*) combined with 5,815 2D class averages from cryoSPARC (**Table 1**), labeled by members of the three participating labs. The combined datasets include 2D averages generated by both RELION and cryoSPARC, effectively training the CNN to recognize and account for platform-specific differences **Fig. 1** . Consequently, CryoSift can generalize its predictions across RELION- and cryoSPARC-generated images. The model outputs labels in STAR file format that users can inspect using the *relion display* function (**Fig. 1B**). CryoSift can be installed as a standalone https://github.com/sstagg/Magellon/tree/main/Sandbox/particle_processor and incor-porated into cryoSPARC user workflows. Alternatively, users can access the online version of the assessor at https://www.cryosift.org, **Fig. S4A**) and can upload either RELION or cryoSPARC-generated 2D class averages to inspect their class quality and optionally contribute their own class labels to refine the CNN model. The latter feature will allow us to continually refine the model with different types of samples and conditions. In this way, the model has the potential to become more accurate and generalizable over time.

Overall, our re-designed 2D class ranker CryoSift offers several advances over our earlier 2DAssess tool. 2DAssess offered no GUI-support, while CryoSift offers an easy-to-use webserver. Its improved CNN architecture supports adaptive average pooling, which is not included in either RELION’s Class ranker or 2Dassess, excluding variable image size as input. The fixed 4-category labeling of 2DAssess cannot pick up on minor differences in 2D class quality, which the continuous labeling within CryoSift can. Adopting RELION’s use of metadata contributed to variations in the final class scoring, adding to CryoSift’s overall improved accuracy.

### Automation with cryosparc-tools

We integrated output from CryoSift as a filter for automatic selection of 2D averages using the *cryosparc-tools* Python library (https://github.com/cryoem-uoft/cryosparc-tools). Even well-aligning 2D classes with visible secondary structure features retain heterogeneity. Less frequent views or low signal-to-noise ratio images may be attracted through an “attractor effect,” ^12^ and iterative 2D classification has proven effective in mitigating this phenomenon. Interfacing with cryoSPARC enables automated and standardized selection of 2D classes through iterative selection and classification of well-aligning classes, while noise and poorly aligning particles are discarded. User-defined thresholds guide the sorting of good and bad particles.

Our *cryosparc-tools* implementation of CryoSift currently iterates 2D classification and labeling over particles with labeling scores between 2.5 and 4.5. In each iteration, 70 % of particles with scores better than or equal to 2.5 are excluded from further iterations, while 30% of particles with scores better than or equal to 2.5 are accepted for the iterative classification. Particles worse or equal to a CryoSift score of 4.5 are discarded. In the following iterations, particles with scores better than 4.5 are pooled, while particles with worse labels are rejected. The number of iterations of processing depends on the extraction box sizes as an indirect measure of particle size. Consequently, box sizes larger than 200 px iterate five times, particles with box sizes between 200-300 px iterate three times, while larger box sizes only iterate twice through the classification and labeling (**Fig. 2**). Subsequently, all particles passing the 4.5 cutoff are pooled and subjected to final 2D classification, then split into three batches with score thresholds of 2.5, 3.5, and 4.5. These batches undergo re-extraction without binning and are subjected to ab-initio reconstruction. The integrated CryoSift framework is illustrated in **Fig. 2**.

**Figure 2.**
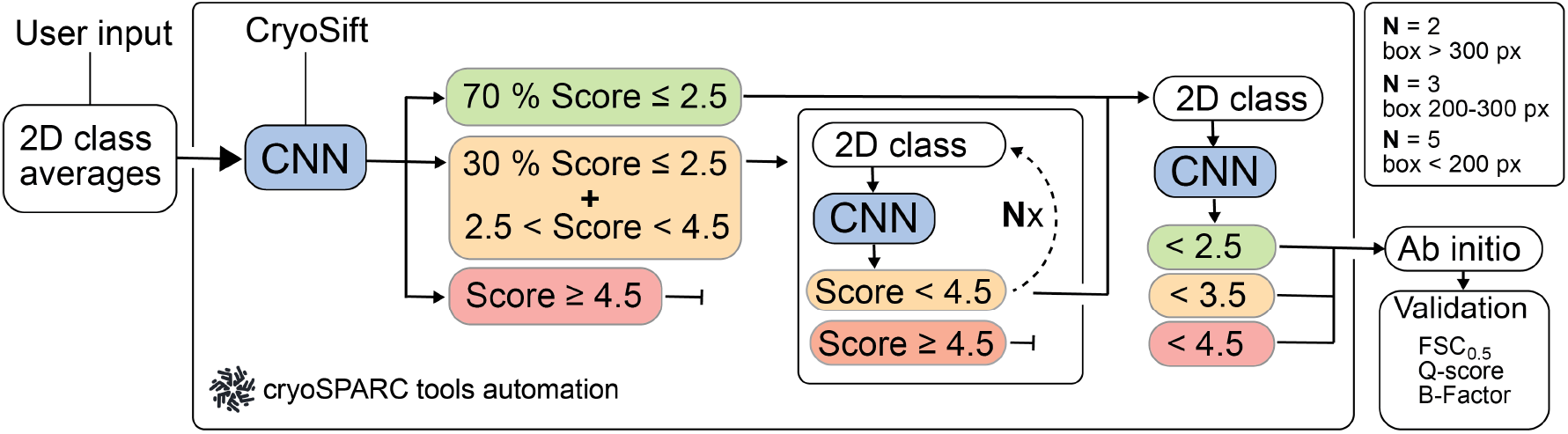
Scheme for the automated workflow for iterative 2D classification using cryosparc-tools. CryoSift generates quality labels from user-provided 2D projections and is forwarded to the CS-tools API. A user-provided cutoff score defines the Select / Reject list for the 2D Select job in cryoSPARC and iterates through 2D classification, class quality labeling and selection for N rounds: N=2 for boxes *>* 300 px, N=3 for boxes between 200 to 300 px and N=5 for boxes *<* 200 px. All particles passing the cutoff criterion are pooled and subjected to an ab initio reconstruction. The reconstruction quality is assessed from the Fourier shell correlation (FSC) to a simulated prior (FSC cutoff at 0.5). The atom inclusion is calculated using a corresponding model file and reported as the mean overall Q-score.

### Validation and Testing of CryoSift

For validation of our CNN model, 3,220 class averages, or 10 % of the original dataset was used. This subset was excluded from training to maintain independence of validation. Our model was trained for 200 epochs in PyTorch using an Adam optimizer ^9^ for the mean-squared error of the predicted score against the labeled score. A learning rate and weight decay of 0.0001 with batch sizes of 32 have been identified as optimal through trial and error, and achieved about 0.0055 MSE loss on the validation data, with insignificant improvement after 60 epochs (**Fig. 1C**). The mean-square error (MSE) between true (labeled) and predicted scores was calculated after conversion to a bin-range from 0-1 to match the 2D class ranker labels from RELION5. An average MSE of 0.0039 was calculated. While high bins and low bins showed the least disagreement with the labels, higher discrepancy was observed for the midrange scores (bins around 0.5) (**Fig. 1C**).

To test our CryoSift, ten datasets were selected, covering a broad range of molecular weights (**Fig. 3A**), oligomeric states, and commonly used microscope-setups (**Table 2**). After standardized user-based preprocessing (outlined in the method section), all datasets were subjected to our unsupervised iterative 2D classification using the cryosparc-tools API as shown in **Fig. 2**. To validate the impact of particle selection based on their CryoSift score, we grouped the particles in three clusters with scores lower or equal 2.5, lower or equal 3.5, and lower or equal 4.5. Since refinements rely on particle weighting schemes ^13^, we generated initial reconstructions using the “Ab-initio reconstruction” job-type in cryoSPARC without symmetry. Recognizing that using stochastic gradient descent with randomized initial seeds can generate biased volumes, we calculated a global resolution estimate of the ab initio reconstructions against simulated data. The simulated data was generated using the corresponding atomic models (**Table 2**), filtered to 8 Å . The FSC was calculated using EMAN (v.2.12) ^14^. Additionally, we calculated the Q-score as a measure of atom inclusion using ChimeraX tool-shed (https://github.com/tristanic/chimerax-qscore) and used cryoSPARC’s ResLog analysis job to calculate the B-factor as an overall estimate for data quality and impact of the imaging conditions and chosen processing steps, focusing on the applied 2D accessor cutoffs ^15^.

**Figure 3.**
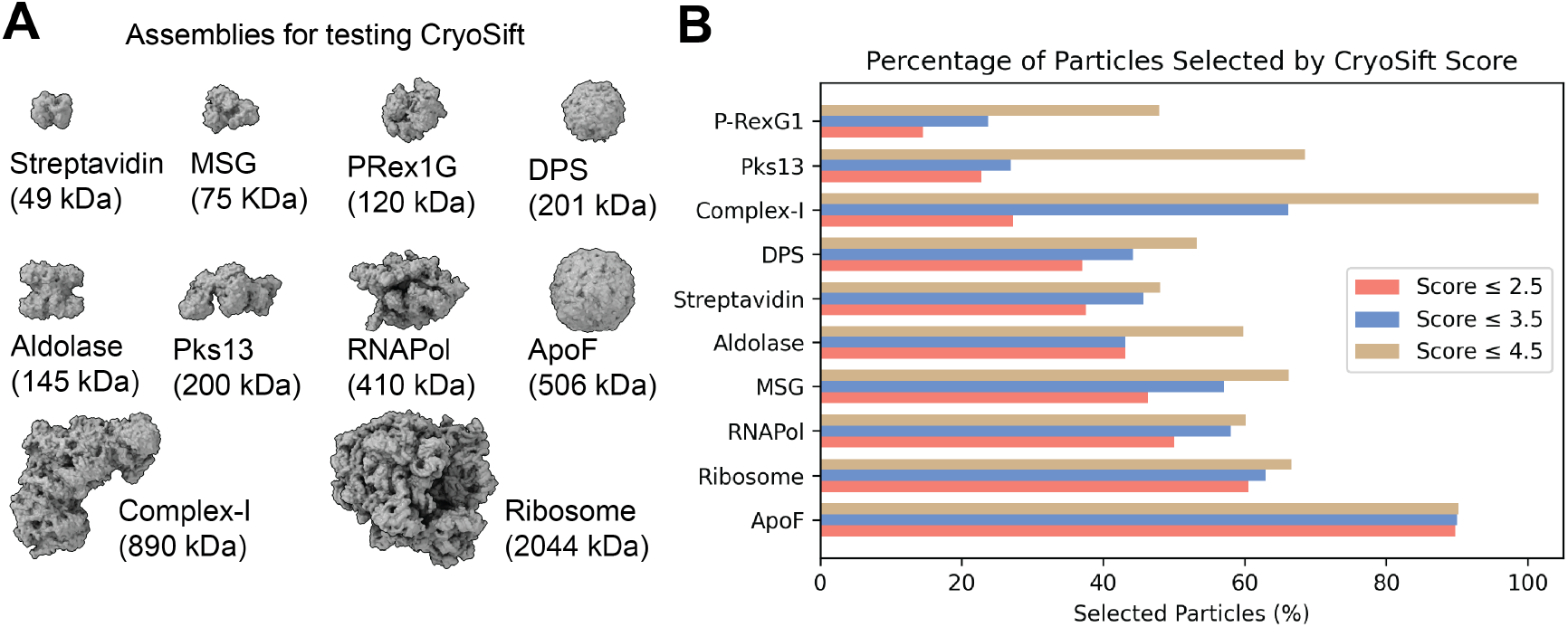
Particle statistics of the automated, unsupervised iterative 2D classification workflow. (**A**) Tested proteins or protein-complexes, ordered by increasing molecular weight. (**B**) Fraction of selected particles according to the CryoSift scores of ≤ 2.5 (coral), ≤ 3.5 (blue) and ≤ 4.5 (tan), ordered by fraction of selected particles.

The fraction of selected particles greatly varied across the chosen datasets (**Fig. 3B**). The accepted particles periteration are shown in supplementary **Fig. S6**, **S7**. It should be noted that the fraction of accepted particles is affected by the quality of detecting particles and generally improves with increasing signal-to-noise (S/N). Additionally, larger particles (ribosomes) have fewer false-positive picks compared to smaller particles (streptavidin). In our processing workflow, particle picks were not subjected to particle inspection and curation to highlight the inherent false-positive picking in noisy datasets.

The mass-to-volume ratio impacts 2D class qualities, as evidenced by the fraction of accepted particles (**Fig. 3B**). ApoF on graphene shows the highest fraction of accepted particles with minor differences between the CryoSift cutoff thresholds. A strong difference between accepted particles across the CryoSift score was present for the membrane protein Complex-I, potentially caused by detergent micelle 2D averages. For the P-RexG1 dataset, only about half of the particles are selected and only about a quarter of the particles pass the 3.5 CryoSift cutoff. For most datasets, the difference in accepted particles between the 4.5 to 3.5 CryoSift cutoff is greater than the difference between 3.5 and 2.5, indicating an optimum for retaining rare views while discarding poorly aligning particles. To validate the quality of the selected particles using our CryoSift score, we generated reconstructions for each of the CryoSift cutoffs. Since cryoSPARC integrates particle weighing in refinement jobs, we used the *ab initio* reconstruction job type without enforcing symmetry, as outlined in the method section. All reconstructed volumes are ordered by increasing resolution and color-coded by CryoSift cutoff scores in **Fig. 3E**.

To validate the quality of the reconstructions, we calculated the FSC to simulated data using atomic coordinates from **Table 2**. Specifically, the FSC at threshold 0.5 was used to define the resolution of the reconstruction and is plotted against the selected CryoSift cutoffs in **Fig. 4A**. Plotting the resolution differential between the 4.5 and 3.5 assess cutoff **Fig. 4B** indicates a negligible impact for several selected datasets, but a substantial resolution improvement for some datasets (e.g. Aldolase, P-RexG1). Additionally, we used the Q-score as an indicator of map quality and interpretability. The mean all-atom Q-score over the CryoSift scores is shown in **Fig. 4C** and the Q-score change between cutoffs 4.5 and 3.5 is visualized in **Fig. 4D**. The map-model fit shows improvements for many datasets, but ApoF and MSG show a loss of the map’s interpretability when applying a stricter 2D cutoff. Both selected validation criteria indicate that curating 2D classes can help improve the quality of 3D reconstructions. While global resolution estimation using the FSC offers an easy comparison to the ground-truth, local map interpretability reported using the Q-score improves when removing low-quality particles, which might carry radiation damage, air-water interface denaturation, or have been collected from areas of low S/N on the grid (thick ice, non-vitreous ice).

**Figure 4.**
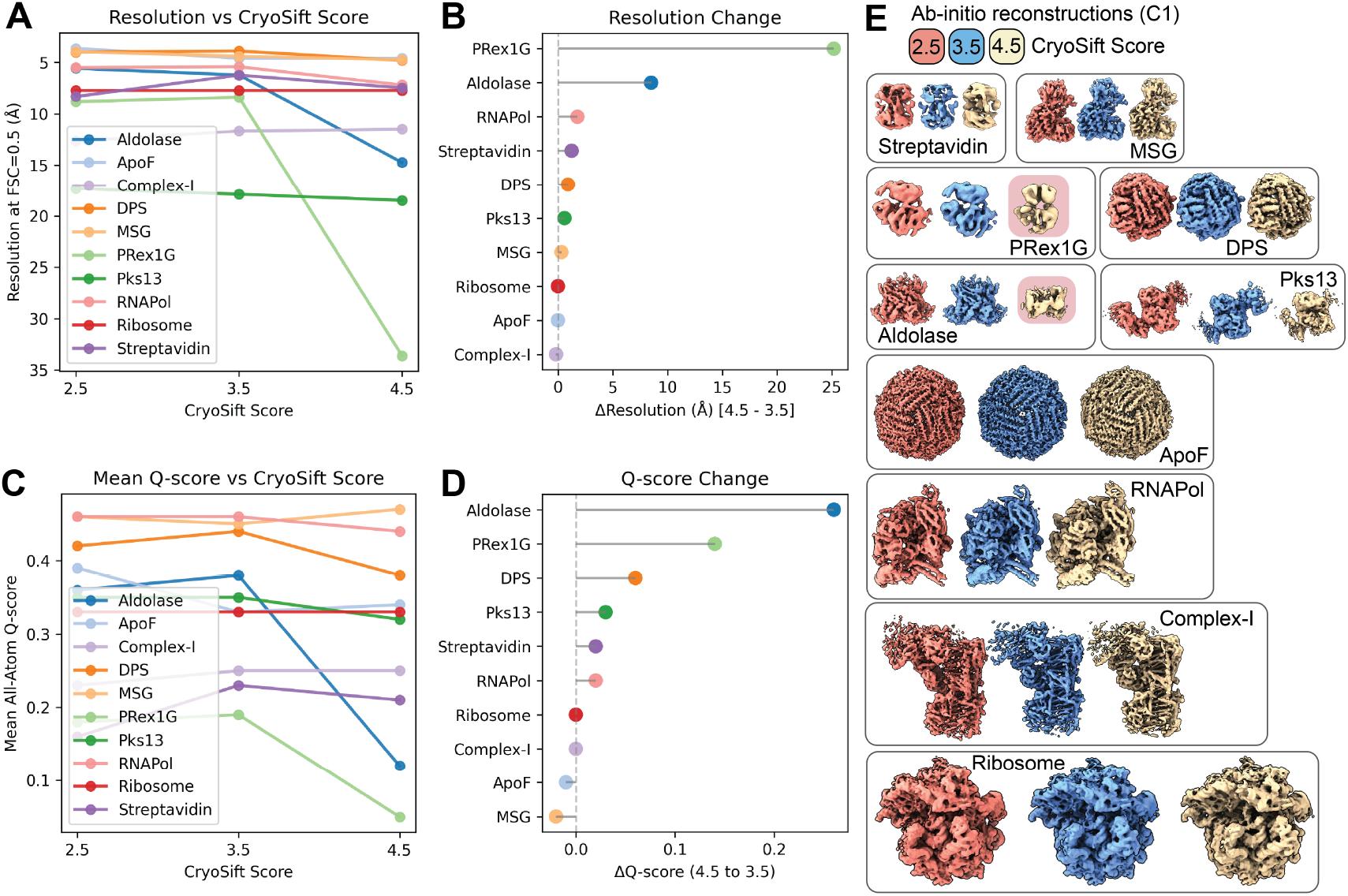
Validation of the reconstructions from the unsupervised iterative 2D classification workflow. (**A**) Resolution of ab-initio reconstructions across three different CryoSift scores using the FSC0.5 against simulated volumes. (**B**) Resolution improvement from CryoSift score 4.5 to 3.5, showing the removal of unsuitable particles. (**C**) All-atom inclusion into ab-initio reconstructions across three different CryoSift scores using the mean all-atom Q-score. (**D**) All-atom inclusion improvement from CryoSift score 4.5 to 3.5. (**D**) Ab-initio reconstructions used for (**A-D**) color-coded by the CryoSift scores. (E) Ab-initio reconstructions were calculated without imposing symmetry (C1) and color-coded by their respective CryoSift score 2.5 (coral), 3.5 (blue) and 4.5 (tan) and ordered by increasing molecular weight. Failed reconstructions for Aldolase and PRex1G highlighted in red.

Degrading resolution and reduction in map interpretability upon removal of particles with stricter labels (cutoff 2.5) is an established observation and correlates the number of particles with a maximum resolution reachable, termed ResLog analysis ^15^. Using cryoSPARC’s implementation, we calculated the ResLog slope which is related to the B-factor for each selected 2D assessor threshold using the ResLog analysis (**Fig. 5**).

**Figure 5.**
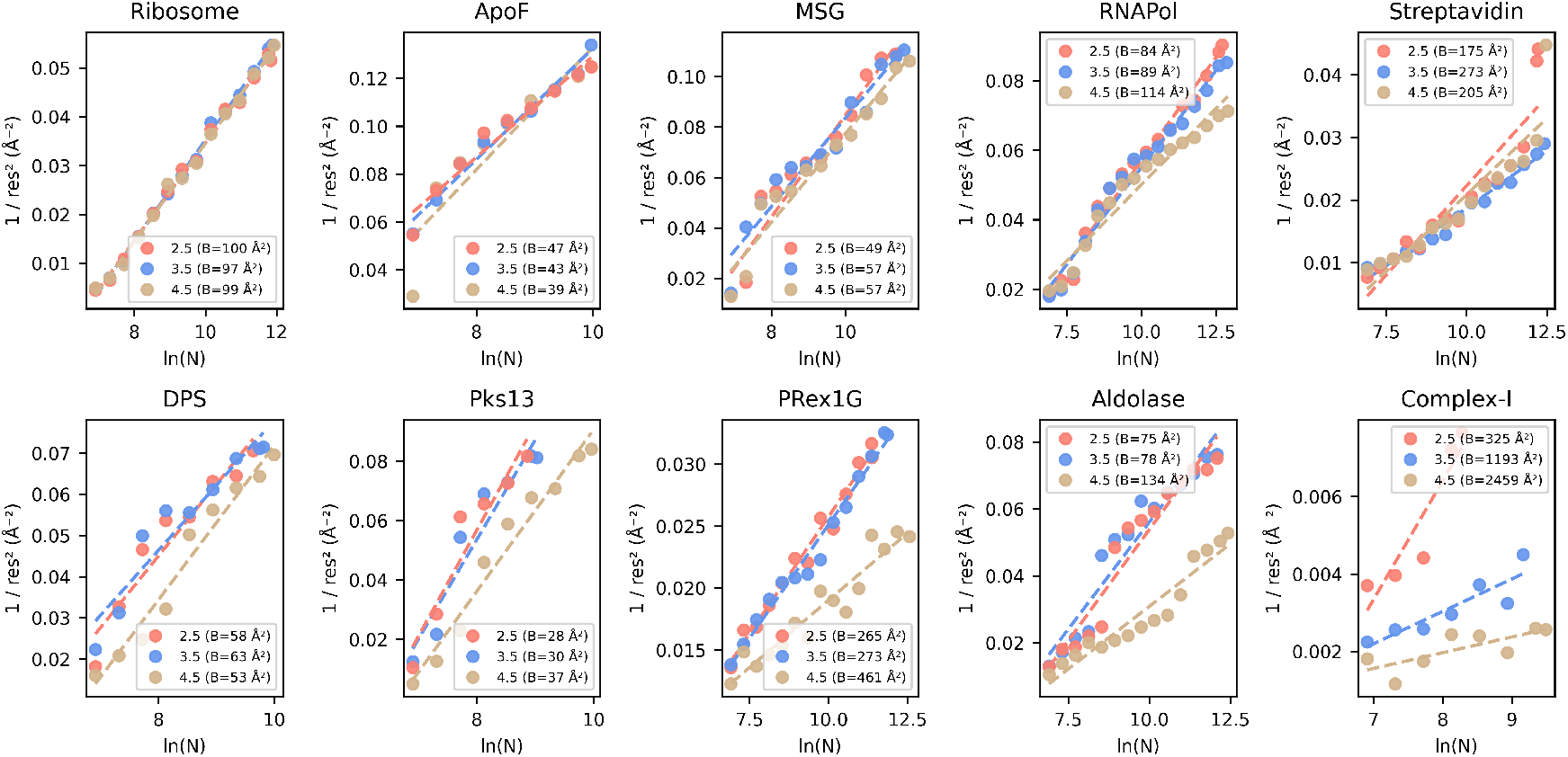
ResLog Analysis of the CryoSift thresholds. Validation of the Reconstruction quality using the ResLog plot (inverse resolution over the logarithm of the number of particles). The slope of the linear regression line, given as B-Factor in Å 2 is an indicator of the data quality (lower, meaning worse quality). Plots were calculated using cryoSPARC’s ResLog Analysis at FSC 0.134. Half-set splits were generated from ab-initio reconstructions using the ‘Homogeneous reconstruction only’ job to preserve the initial angular assignments.

The strongest improvement in map interpretability (Q-score) and overall resolution (FSC 0.5) is also visible in increases in the ResLog slope and thus increased particle quality when comparing the 3.5 with the 4.5 CryoSift cutoffs for Complex-I, Aldolase and PRex-1G. Also, DPS, Pks13 and RNAPol show improvements that match our earlier observations (**Fig. 4**). Almost no changes are visible for the Ribosome and ApoF dataset, since they retain most of their particles when applying the different 2D access cutoffs (**Fig. 3B**).

To test cross-platform usability of CryoSift, we transferred pre-processed particle stacks of Pks13, DPS and Complex-I from cryo-SPARC to RELION5 using *pyem* and generated new 2D class averages using RELION’s 2D classification (VDAM). Subsequent class labeling with our CryoSift server demonstrates its ability to work on both RELION and cryo-SPARC data (**Fig. S4S4**).

Direct comparison of RELION5 processed classes with its Class Ranker and CryoSift using quality-sorted classes demonstrates a quasi-linear correlation across the two class labelers. While the Complex I dataset shows linear agreement across the entire labeling range, Pks13 and DPS show variations in assigned class quality for mid-tier class qualities, indicating a finer grained quality estimation of CryoSift over RELION’s Class Ranker (**Fig. S5**). An observation, consistent with the validation of the CNN (**Fig. 1C**), potentially pointing to diverging class quality assignment by the curators during training.

## Discussion

In this paper, we present a platform-agnostic 2D class average assessor, termed CryoSift, that addresses a key bottleneck in cryo-EM data processing: unsupervised automation across user- and experience-levels with minimal integration efforts. We demonstrate the versatility of CryoSift by integrating it into cryoSPARC using the cryosparc-tools API, creating a fully automated workflow that accommodates users across different experience levels. At its core, the CNN-based CryoSift labels 2D class averages with quality scores, which can be used either as a stand-alone application for assessing RELION or cryoSPARC 2D averages, or as an integrated component within automated processing pipelines (such as cryoSPARC’s *Select 2D* job utilizing the cryoSPARC tools API). This iterative approach to 2D classification, quality labeling, and selection helps promote the inclusion of rare views that would otherwise be discarded in non-iterative procedures ^16^. A systematic evaluation of ten diverse datasets using FSC-based resolution estimates, Q-score atom-inclusion metrics, and B-factor analysis revealed that a quality score cutoff of 3.5 provides an optimal balance between including rare views (lower quality classes) and excluding unsuitable classes or false-positive picks.

While symmetric, high-molecular weight targets (like ApoF and DPS) generate stable 2D classes without meaningful improvements from iterative classification (**Fig. 3B, 4E, S6**), most research-relevant samples suffer from conformational heterogeneity, compositional variability, and preferred orientation effects. Our aldolase dataset exemplifies how iterative classification successfully removes particles affected by partial denaturation and localized damage, as observed in EMD-21492, demonstrating the tool’s value for real-world applications (**Fig. 4E, S6**).

A notable strength of this pipeline is its multi-tiered accessibility. Users new to cryo-EM analyses can upload data to our web server to familiarize themselves with expert-trained quality assessments, then apply default cryosparc-tools settings to their own datasets. Intermediate-level users can adjust parameters for their specific targets and implement parallel processing workflows. Advanced users and developers can contribute to CNN optimization by sharing labeled datasets, extend workflows for more challenging systems (e.g. filaments and membrane proteins) and combine cryosparc-tools with platform conversion utilities for cross-platform automation. We have demonstrated this using *pyem* for file conversion of cryo-SPARC particle stacks to RELION, and performed a direct comparison using three different datasets, which highlight easy cross-platform use of CryoSift with comparable class quality labeling (**Fig. S4, S5**). While this manual approach of class quality labeling using CryoSift for RELION data can be informative, the biggest benefit arises from automation using inbuilt cutoffs and iterative 2D classification with pooling of particles across multiple 2D classification and selection jobs. Consequently, developing a RELION-compatible iterative workflow would result in higher reconstruction quality and reduces the need for user-intervention. In its current state, a conversion and transfer to cryoSPARC with subsequent use of our CryoSift and cryosparc-tools pipeline would enable easy automation of 2D class selection.

We envision CryoSift to serve as a community-driven tool that improves through user contributions. The web server enables users to contribute their own class labels, allowing continuous model refinement with diverse sample types and conditions. This crowdsourced approach has the potential to make the model increasingly accurate and generalizable over time. Further, this tool has relevance in educational purposes, as it provides clear examples of quality differences in 2D class averages, making it valuable for cryo-EM training programs.

We note that several extensions would further enhance the CryoSift’s utility. The substantial fraction of false-positive picks observed in our datasets (**Fig. 3B**) suggests that automatic clustering of picked particles (cryosparc v.4.6+) without requiring user-defined inputs could increase processing speed and improve reconstructions. Beyond quality-based selection, incorporating projection-level information to assess compositional heterogeneity, detect preferred orientation effects, and compare particle distributions to simulated projections would provide valuable additional capabilities. Further, our assessment tool currently focuses on overall class quality rather than distinguishing between different conformers, oligomeric states, or contaminants. While this approach effectively removes poor-quality classes, future development could incorporate more sophisticated classification schemes.

Our integration of CryoSift into cryoSPARC and planned incorporation into the Magellon platform for cryo-EM visualization, management, and processing ^17^ (https://www.magellon.org/) represents a step toward fully automated cryo-EM processing pipelines that maintain user control while reducing the burden of repetitive tasks. We anticipate that this approach has the capacity to accelerate structure determination and increase accessibility of high-quality cryo-EM analyses to the broader scientific community.

## Acknowledgements

Funding for this research was provided by a National Institutes of Health grant R01 GM143805 to SMS, GCL, and MAC, and the German Research Council project number 556478029 to JHS. The authors thank Jean-Christophe Ducom and Charles Bowman at Scripps for computational support.

## Author contributions

The CNN code was written by KH and PH. The cryoSPARC implementation was realized by JHS and AC. The initial draft manuscript was written by JHS and GCL and subsequently edited by all authors.

## Data & software availability

The in-house mouse apoferritin dataset used for testing was deposited as EMD-70880, and aligned dose-weighted micrographs are available on the Electron Microscopy Public Imaging Archive (EMPIAR-12798). CryoSift is available on GitHub (https://github.com/sstagg/Magellon/tree/main/Sandbox/particle_processor), and the CryoSift web server is available at https://www.cryosift.org.

## Supporting Information

**Figure S1.**
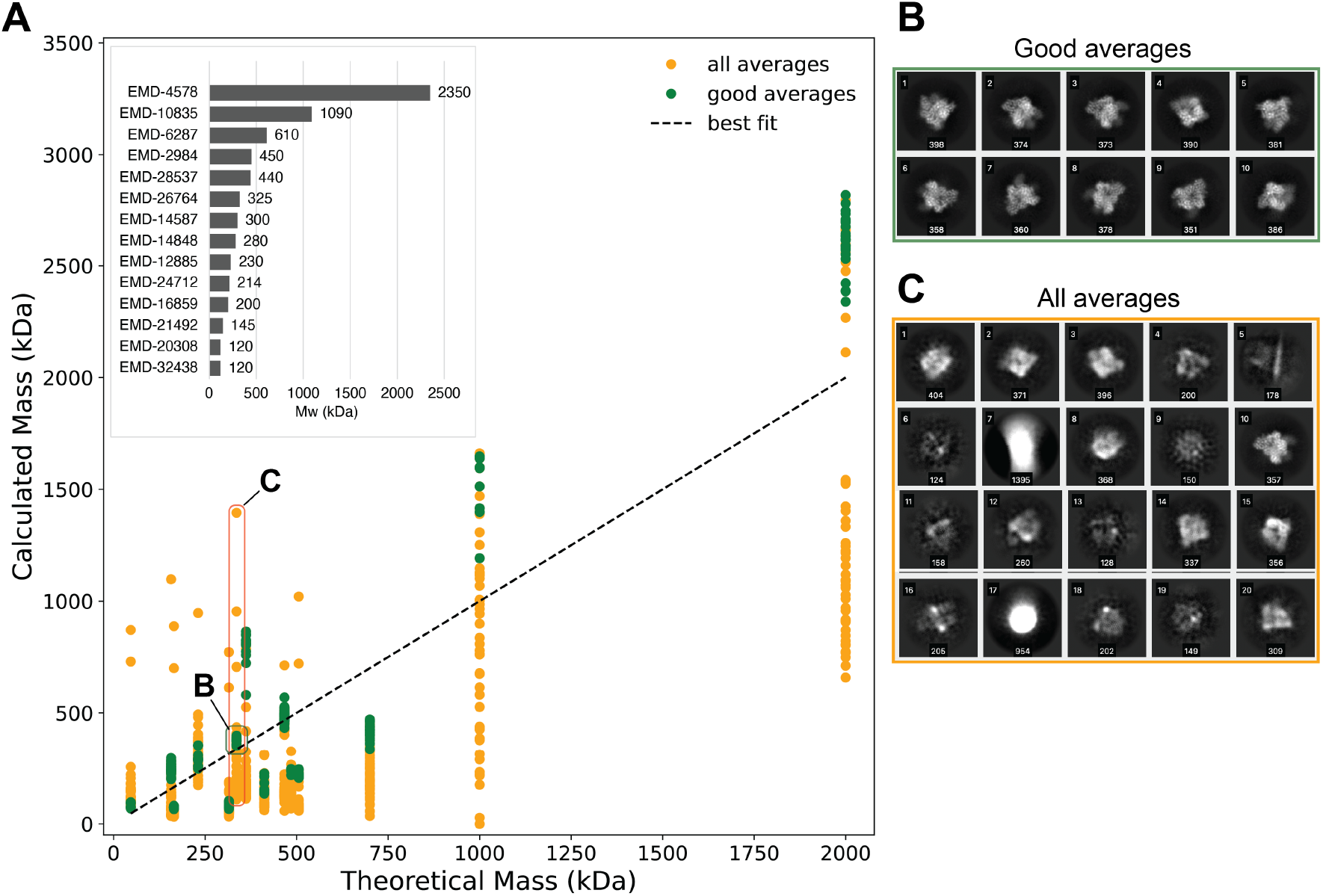
Mass Estimator. (**A**) Correlation of the calculated mass over theoretical mass (in kDa). All averages in orange and good averages in green. Linear fit as dashed lines. Inlet shows the utilized EMD entries sorted by molecular weight. (**B**) Examples of good 2D class averages with calculated mass labels. (**C**) range of all classes in selected example with calculated mass labels.

**Figure S2.**
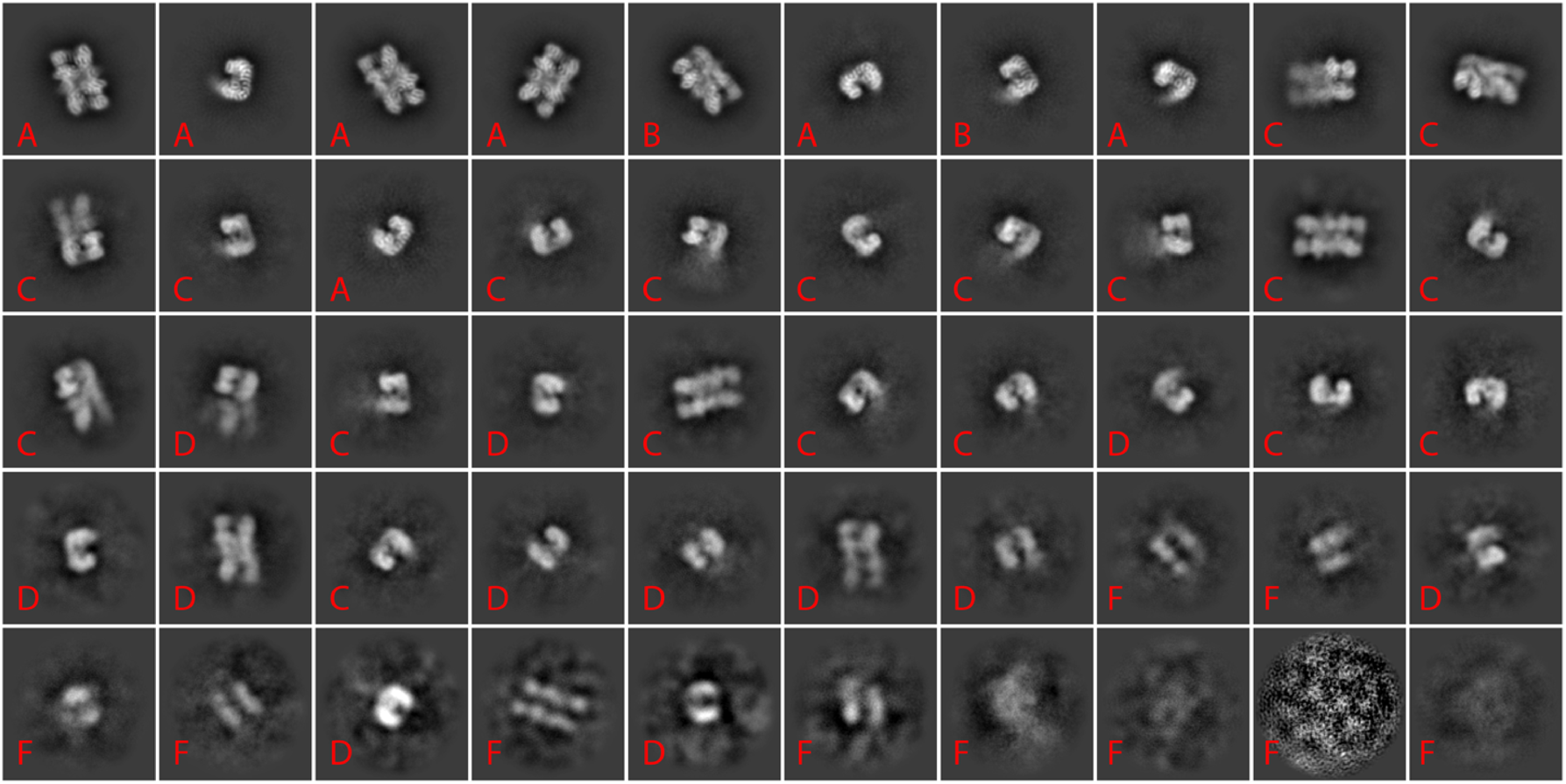
Example labeled class averages included in the grading rubric for assessors.

**Figure S3.**
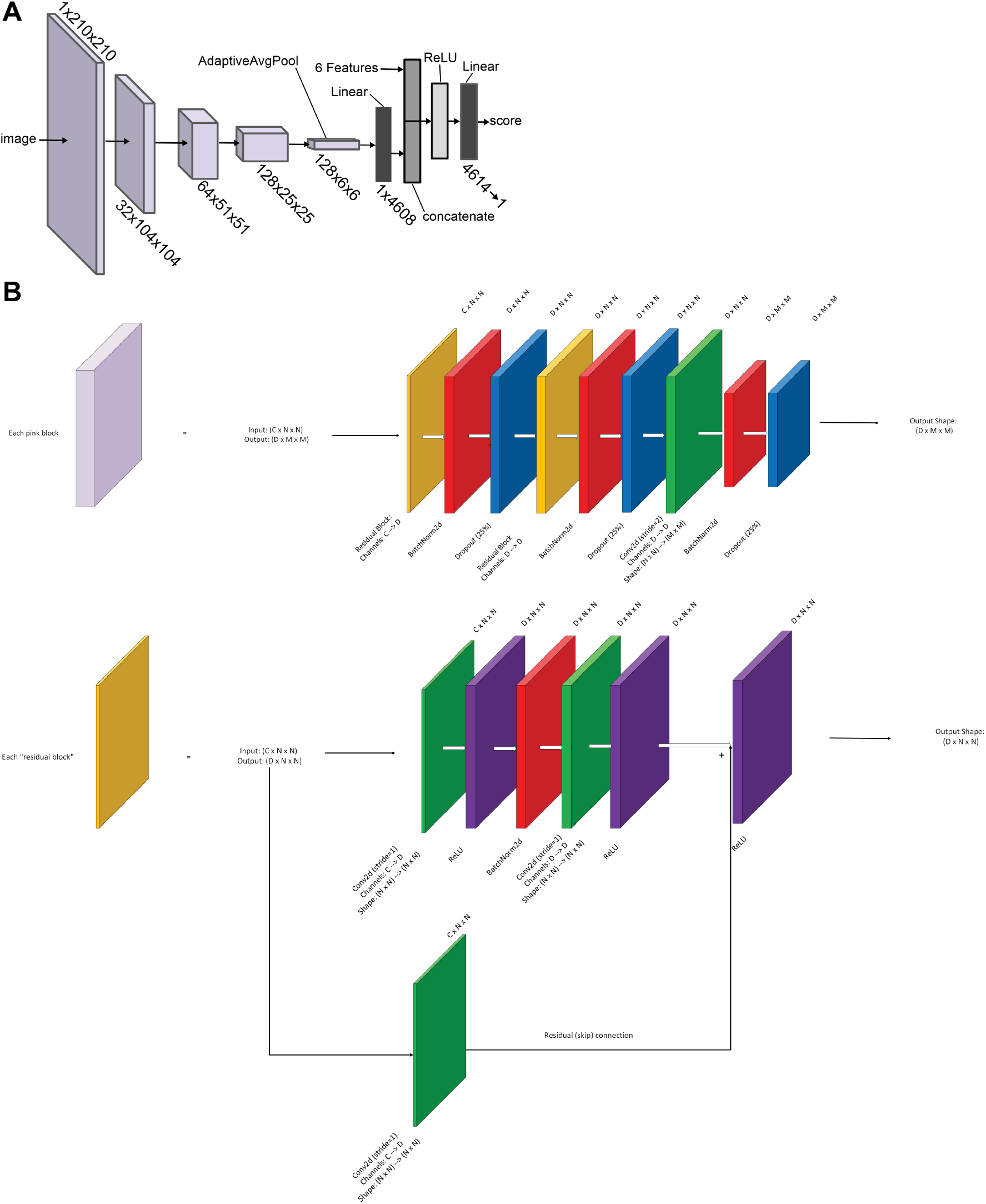
Detailed CNN design architecture. (**A**) Overview of the base layers. (**B**) Detailed layer contents and operations for main layers and each “residual block”.

**Figure S4.**
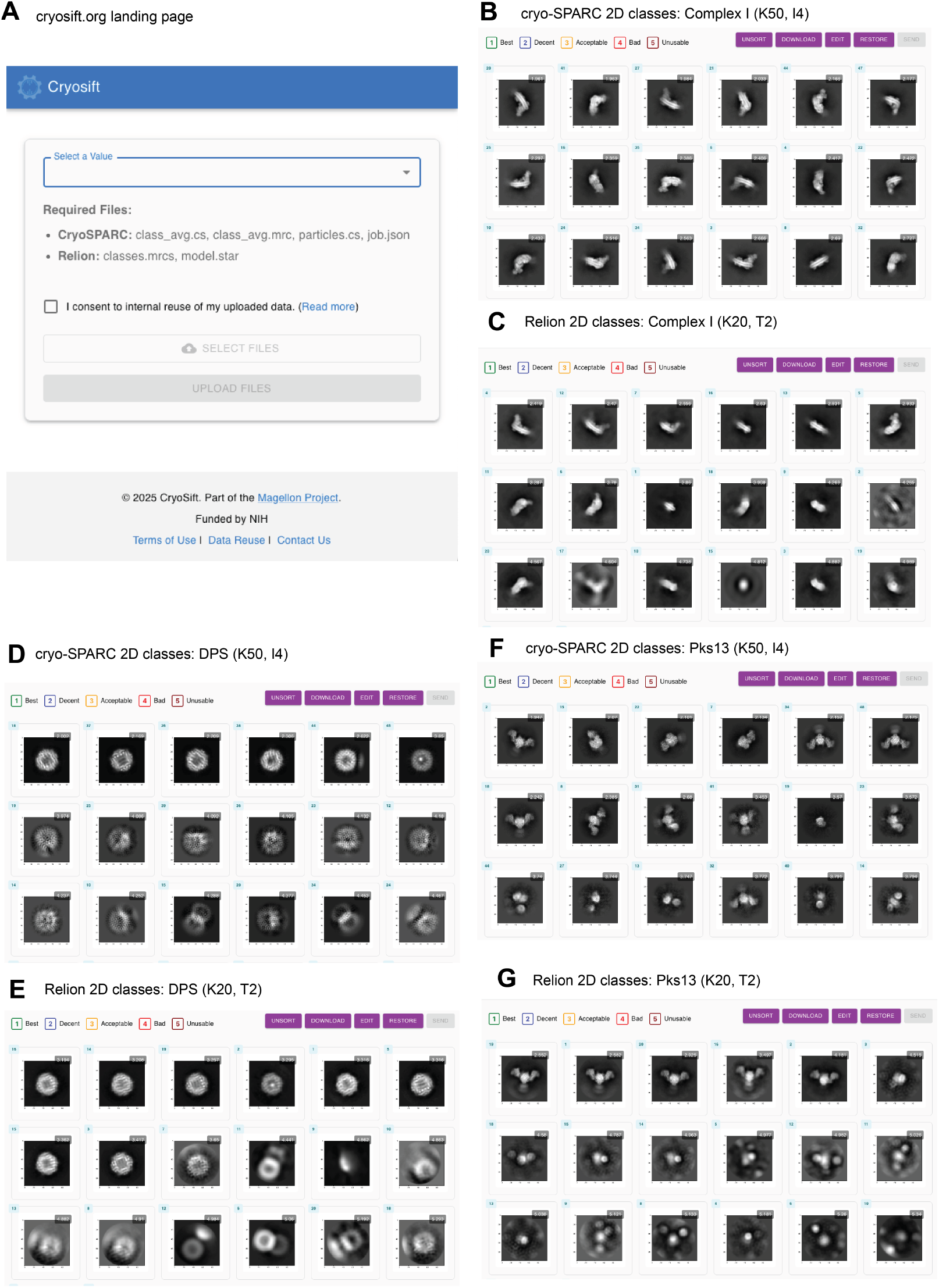
Preview of the CryoSift GUI available at https://www.cryosift.org. (**A**) Landing page. (**B**) Output of sorted Complex-I 2D classes from cryo-SPARC uploads (K50, 50 classes and I4, initial class uncertainty factor 4). (**C**) Output of sorted Complex-I 2D classes from RELION5 (VDAM, K50, T2). RELION re-processing from a re-extracted particle-stack using pyem for file conversion from cryo-SPARC.

**Figure S5.**
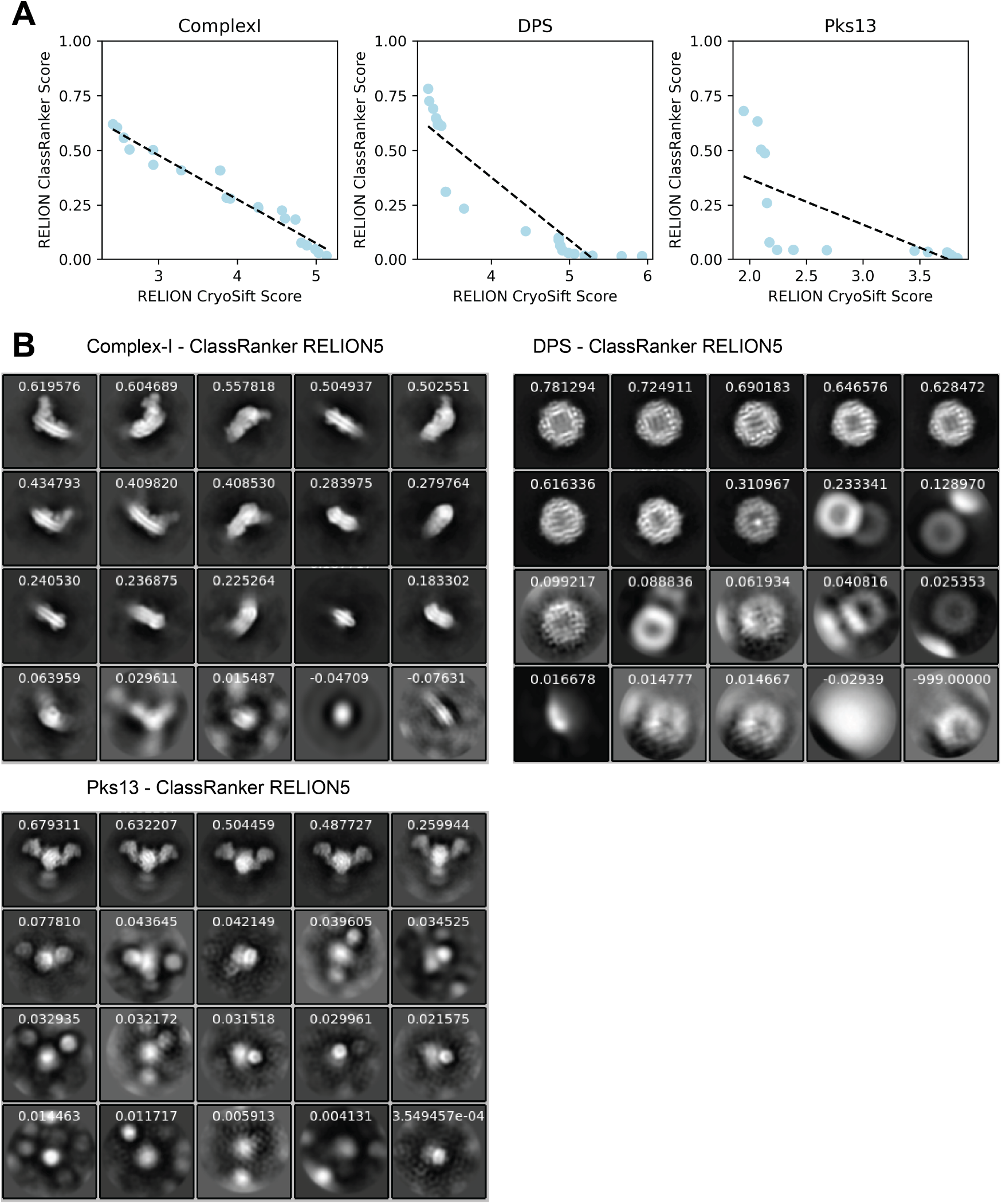
Comparison of labeling 2D classes. (A) Scotter plot and linear fit of RELION 5 processed datasets, showing it’s native ClassRanker scored classes over corresponding CryoSift scored classes. (B) 2D class averages and ClassRanker scores used in (B). 2D averages and CryoSift scores used for (B) shown in Fig. S4.

**Figure S6.**
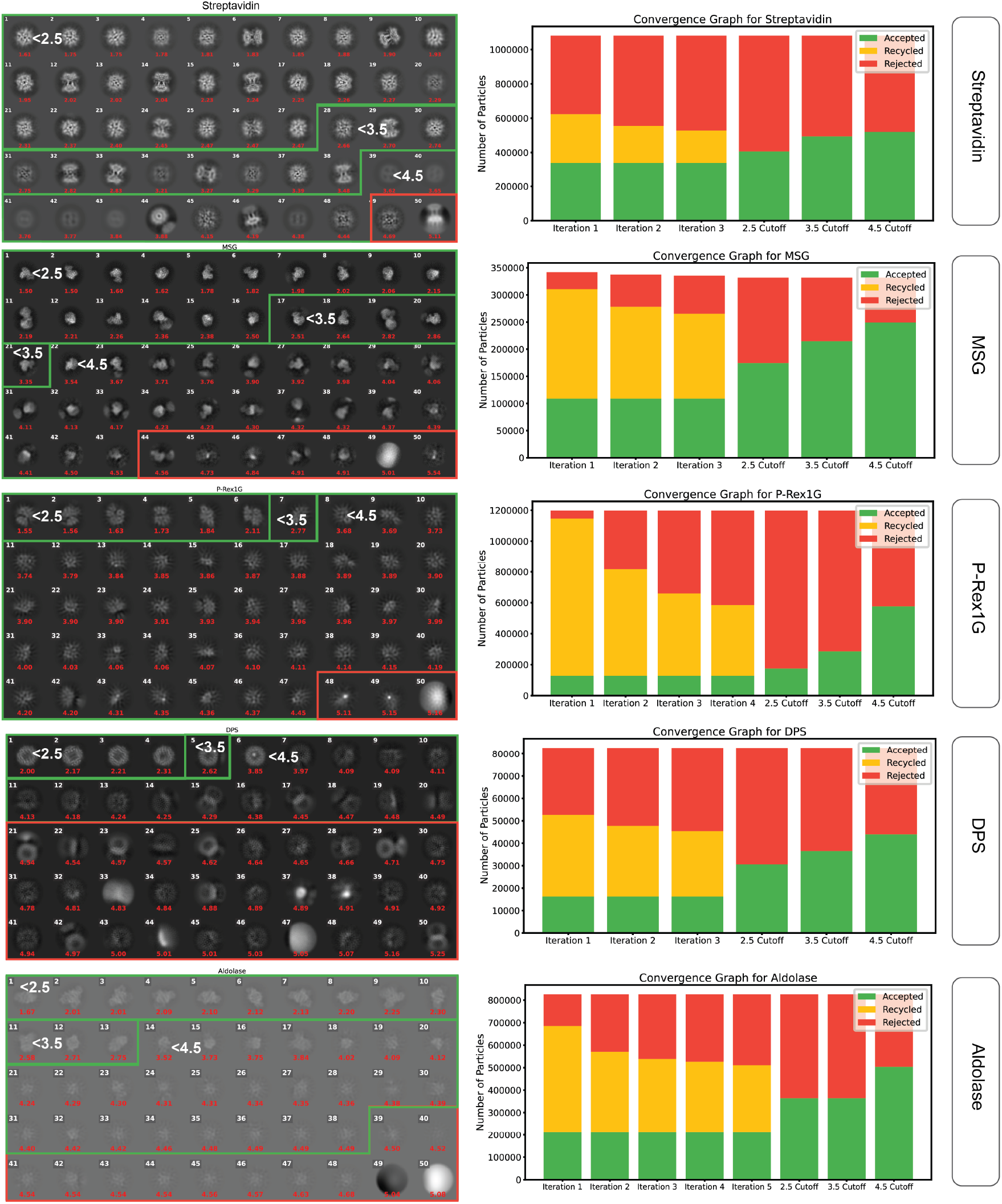
Labeled 2D averages of the last 2D classification for all test-datasets ordered by increasing CryoSift score (red) with associated convergence graphs for all iterations. Projections of matching cutoffs are grouped using green (accepted) and red (rejected) boxes using the respective threshold in white. For the convergence graphs, particle numbers were plotted over iterations. Accepted particles (green), recycled particles (yellow), and rejected particles (red). Additionally, the final number of accepted (green) and rejected (red) particles per CryoSift cutoff score (2.5, 3.5 and 4.5) are depicted.

**Figure S7.**
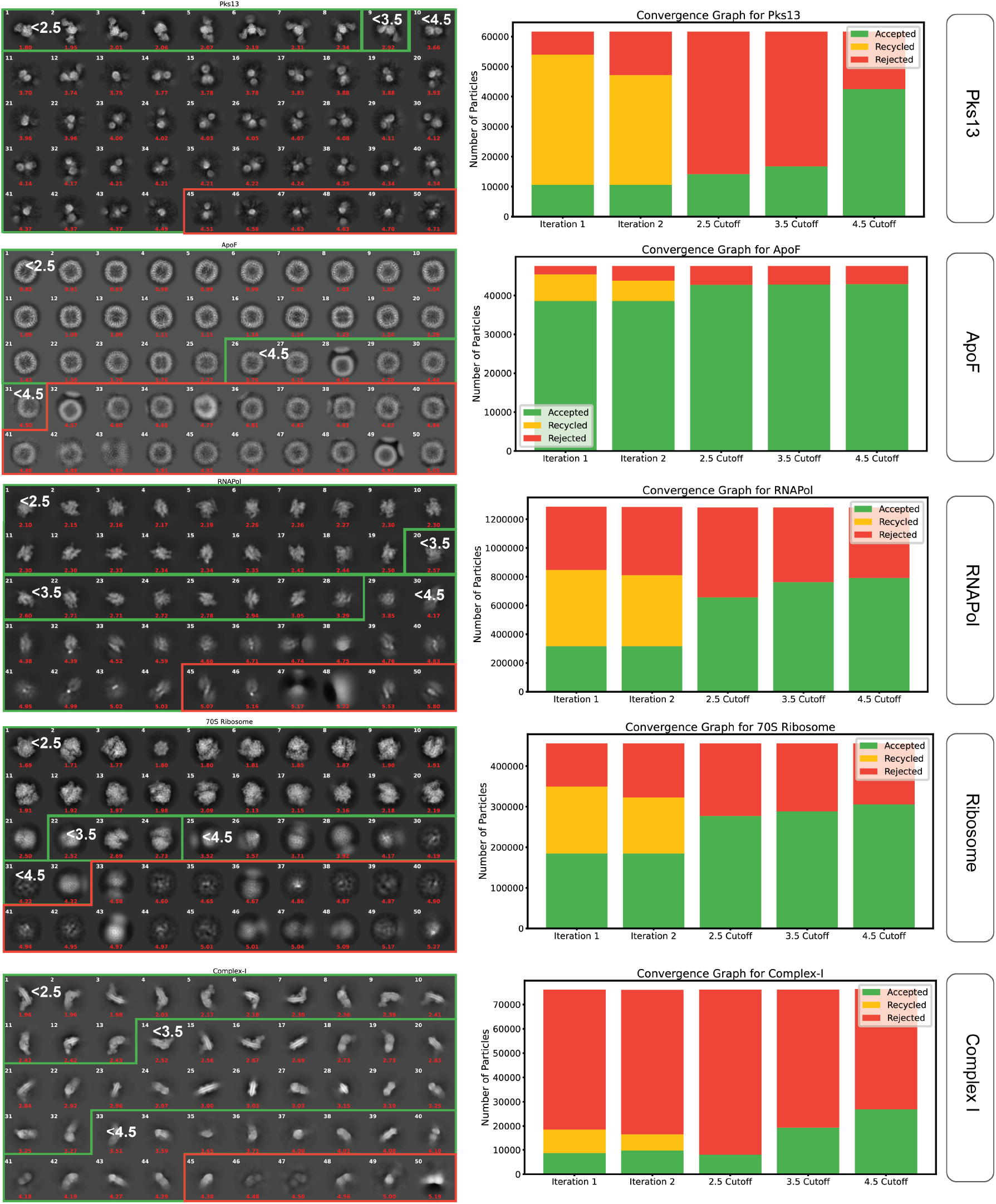
Continued

